# In Silico analysis of PE_PGRS20 (Rv1068c) protein in *Mycobacterium tuberculosis* H37Rv

**DOI:** 10.1101/2021.08.31.458215

**Authors:** Saleem Ahmad, AnupKumar Kesavan

## Abstract

The genetic makeup of *<u>Mycobacterium tuberculosis</u>* reveals the presence of an unknown repeat sequence of PE_PGRS family proteins that are responsible for antigenic variations and many unknown functions that includes necrosis of macrophage and apoptosis. The structure and function of these glycine-rich proteins can be predictable by homology modeling, the Ab-initio method, or by using different tools of bioinformatics. In this study, we selected, PE_PGRS20 (Rv1068c) an unknown PE_PGRS protein. We suggest that the PE_PGRS20 gene is linked with the others genes of the *espACD* operon which are the virulence factors in the *<u>M</u>*. *<u>tuberculosis</u>* H37Rv strain. The genes associated with this protein secretion system can perform the synthesis of a special type of fatty acid known as phthioceroldimycocerates (PDIM).

Docking with different anti TB drugs shows binding with PE_PGRS20 protein which suggests that the target protein may involve in the drug resistance.

## Introduction

The decrypting of the *<u>M</u>*. *<u>tuberculosis</u>* genome shows the existence of a unique PE_PGRS gene family encoding proteins consisting of two major domains: a PE domain linked to a PGRS domain, which contains numerous glycine and alanine-rich repeats^1^. There is much interest in this gene family since, to date, its members have been found only in the genomes of pathogenic Mycobacteria. Understanding the function, localization, regulation, and host response to PE_PGRS proteins is vital for determining what role these proteins may play in mycobacterial pathogenesis. Proof indicates that some PE_PGRS proteins can be specifically expressed in granulomas, where they may influence infectivity of host cells^2,3^, and can induce host cell necrosis^4^ or apoptosis^5,6^. Regulatory mechanisms affecting PE_PGRS gene expression are unknown, although various PE_PGRS proteins are differentially expressed by *M. tuberculosis* both in vivo^4^and in vitro ^7,8^.

Despite its significance, relatively few in-depth investigations are available. The origin and evolution of this family remain a mystery.^9^

In this study, we choose unknown PE_PGRS20 of *M. tuberculosis* H37Rv strain to solve the structure by using homology modeling and ab-initio modeling method. Different databases were used to solve the structure of PE_PGRS20 protein and prediction of the function was done by molecular docking method.

## Materials and Methods

### Structure prediction

The amino acid sequence of the target protein, PE_PGRS20, was obtained from the Mycobrowser (https://mycobrowser.epfl.ch/genes/Rv1068c). SWISS-dock, Phyre2, and Modeller were used to identify the structure As the result provided by these servers gave only 17% of the total structure (template: 5xfs.1.A), we used the ITASSER server to predict the whole structure of PE_PGRS20 protein. ITASSER gave five structures based on the C-Score (https://zhanggroup.org/I-TASSER/).^10^ The structure with the lowest C-Score had been chosen for further docking studies. The final structure was refined by ModRefiner (https://zhanglab.dcmb.med.umich.edu/ModRefiner/).^11^ Structure validation is carried out by PROCHECK server (saves v6.0).^12^ Protein sequence fetched from mycobrowser:

>Mycobacterium tuberculosis H37Rv|Rv1068c|PE_PGRS20

MSYMIAVPDMLSSAAGDLASIGSSINASTRAAAAATTRLLPAAADEVSAHIAALFSGHGEGYQAIARQMAAFHDQFTLALTSSAGAYASAEATNVEQQVLGLINAPTQALLGRPLIGNGADGTAANPNGGAGGLLYGNGGNGFSQTTAGLTGGTGGSAGLIGNGGNGGAGGAGANGGAGGNGGWLYGSGGNGGAGGAGPAGAIGAPGVAGGAGGAGGTAGLFGNGGAGGAGGAGGAGGRGGDGGSAGWLSGNGGDAGTGGGGGNAGNGGNGGSAGWLSGNGGTGGGGGTAGAGGQGGNGNSGIDPGNGGQGADTGNAGNGGHGGSAAKLFGDGGAGGAGGMGSTGGTGGGGGFGGGTGGNGGNGHAGGAGGSGGTAGLLGSGGSGGTGGDGGNGGLGAGSGAKGNGGNGGDGGKGGDAQLIGNGGNGGNGGKGGTGLMPGINGTGGAGGSRGQISGNPGTPGQ

### Docking studies

The docking studies were carried out by autodockvina and Easydockvina tools.^13^ The 3D structures of various ligands were downloaded from PubChem (https://pubchem.ncbi.nlm.nih.gov). The 3D structure of the model of the protein PE_PGRS20 was downloaded from the ITASSER server. The blind docking method was used to identify the different active sites. The active site was modeled in PYMOL software and interaction was observed in LigPlus software.^14^

### Function Prediction

The string pathway analysis (https://string-db.org/) was used to predict the function of the target protein. The PE_PGRS20 amino acid sequence was downloaded from the Mycobrowser database and submitted to the String database. Docking study and String analysis together were able to predict the function of protein PE_PGRS20. The motif sites were predicted by the Myhits motif scan (https://myhits.sib.swiss/cgi-bin/motif_scan/). The function of PE_PGRS20 protein was predicted by String Database and Gene Ontology (quickGO).

### Molecular Dynamic Simulations

Molecular dynamic simulation was used to observe the stability of modelled structure. For molecular dynamic simulation Gromacs 5.1.2 was used with the AMBER03 (nucleic, amber94) forcefield and TIP3P water model at standard temperature and pressure. The energy plots was observed with Grace graph plotting tool. PyMol was used to observe the changes occur in the various residues of protein including active site residues.

#### Salt Bridge Analysis

Salt bridge analysis was carried by using trajectory files of MD simulation of protein with the help of VMD software. The graph is plot on grace, The salt bridges contribute to the stability and folding unfolding of protein.

## Results

### 1. Tertiary structure prediction and its validation

The final structure of PE_PGRS20 was synthesized by using template of apoptosome of *D. melanogaster* (PDB ID: 4V4L). ITASSER server was used for structure prediction. It used ab initio method of structure prediction of protein.

For structure validation, the protein structure was refined by ModRefiner server and submitted to PROCHECK server (SAVES v6.0). Ramachandran plot was used for structure validation and 75.0% residues are in favored regions. Errat analysis shows that the structure has 71.9 % overall quality factor and verify3D confirms that the protein structure has 83.59% residues have 3D-1D scored >=0.2. (also see Fig.1 A, B and C)

**Figure.1:**
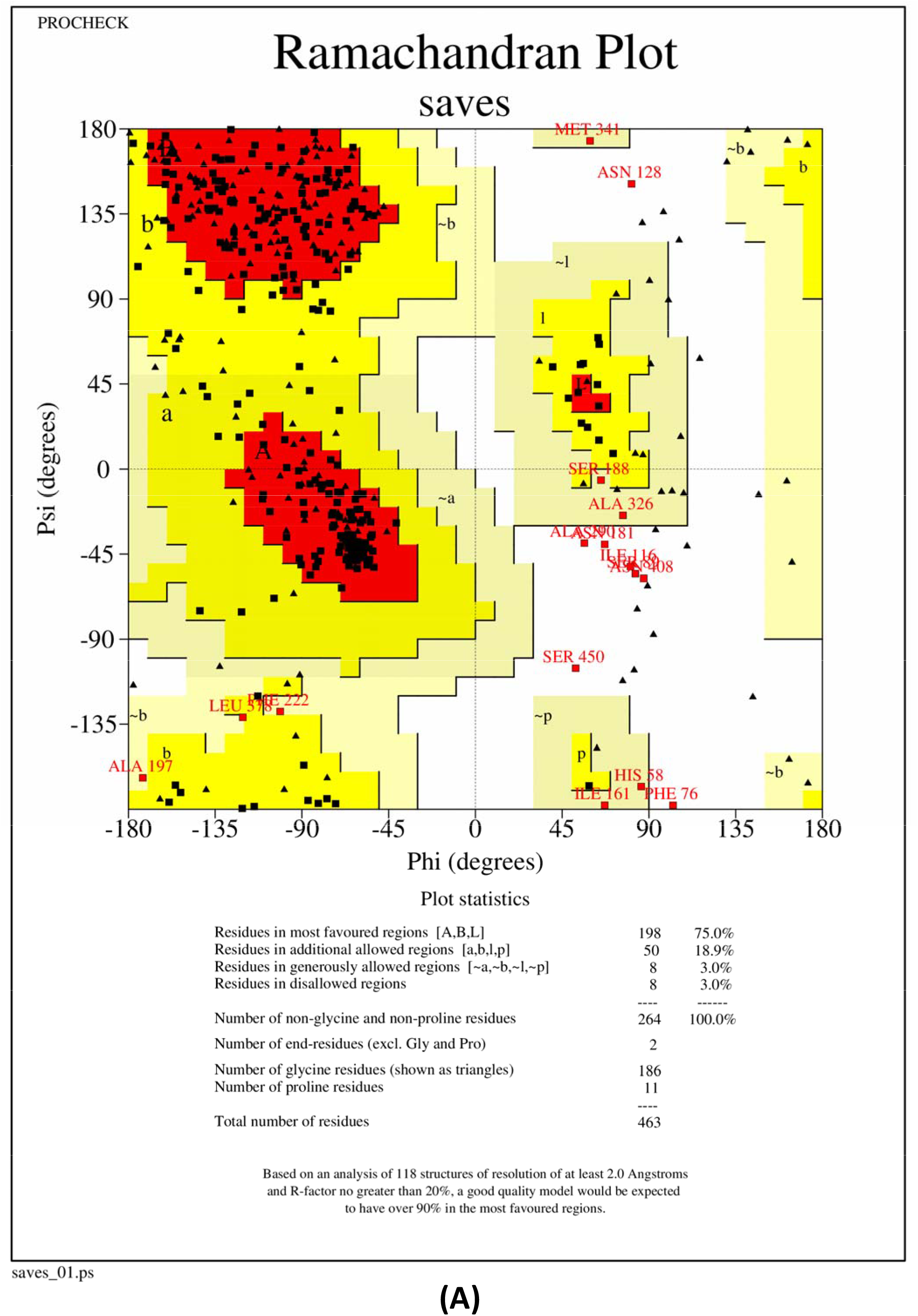
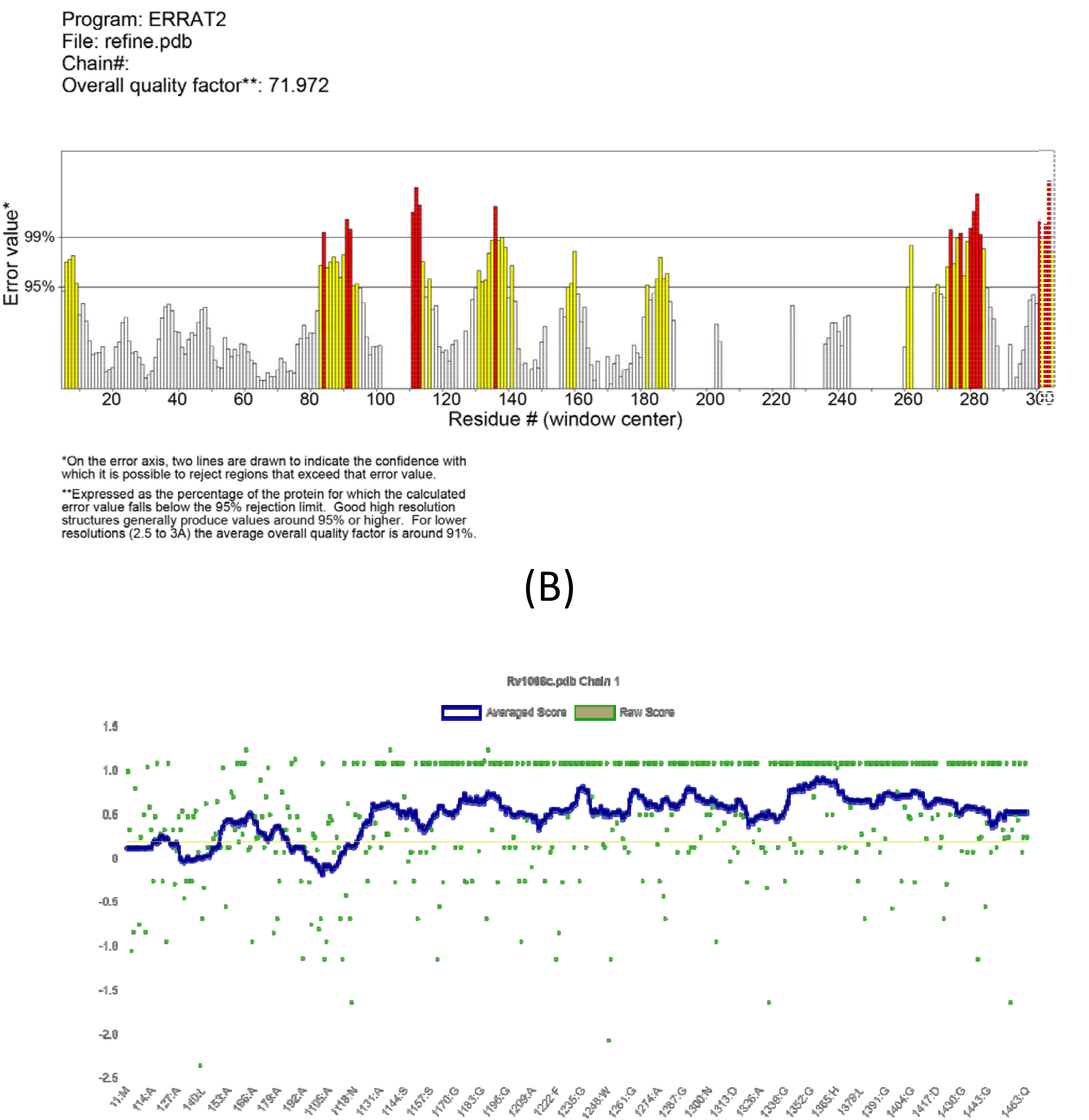
**(A)** Ramachandran Plot Analysis of structure of PE_PGRS20 protein. **(B)** Errat analysis showing overall quality factor of protein structure. **(C)** Verify3D showing the percentage of average score of each residue.

### 2. 3D structure shows the presence of parallel beta-rolls and calcium binding motif

The 3D structure downloaded from ITASSER server showed the presence of parallel beta roll and calcium-binding motif. The calcium binding motif was identified by protein ligand complex binding.^15^The PE_PGRS has conserved N terminal PE domain, which shows high similarity to PE family members and variable C terminal PGRS domain. PGRS domain contains repeats of Gly-Gly-Ala and Gly-Gly-X, where X can be any amino acid.^16^Calcium is very important molecule which is used in many signaling pathways whether it’s a prokaryotic pathway or eukaryotic cellular pathway. PE_PGRS genes containing nona-domain can bind with Ca^2+^ and initiate infection in macrophage through TLR2 (toll-like receptor). This binding modifies IL-10, an anti-inflammatory interleukin, signaling.^17^(see Fig 2.A and 2.B)

**Figure 2.1.**
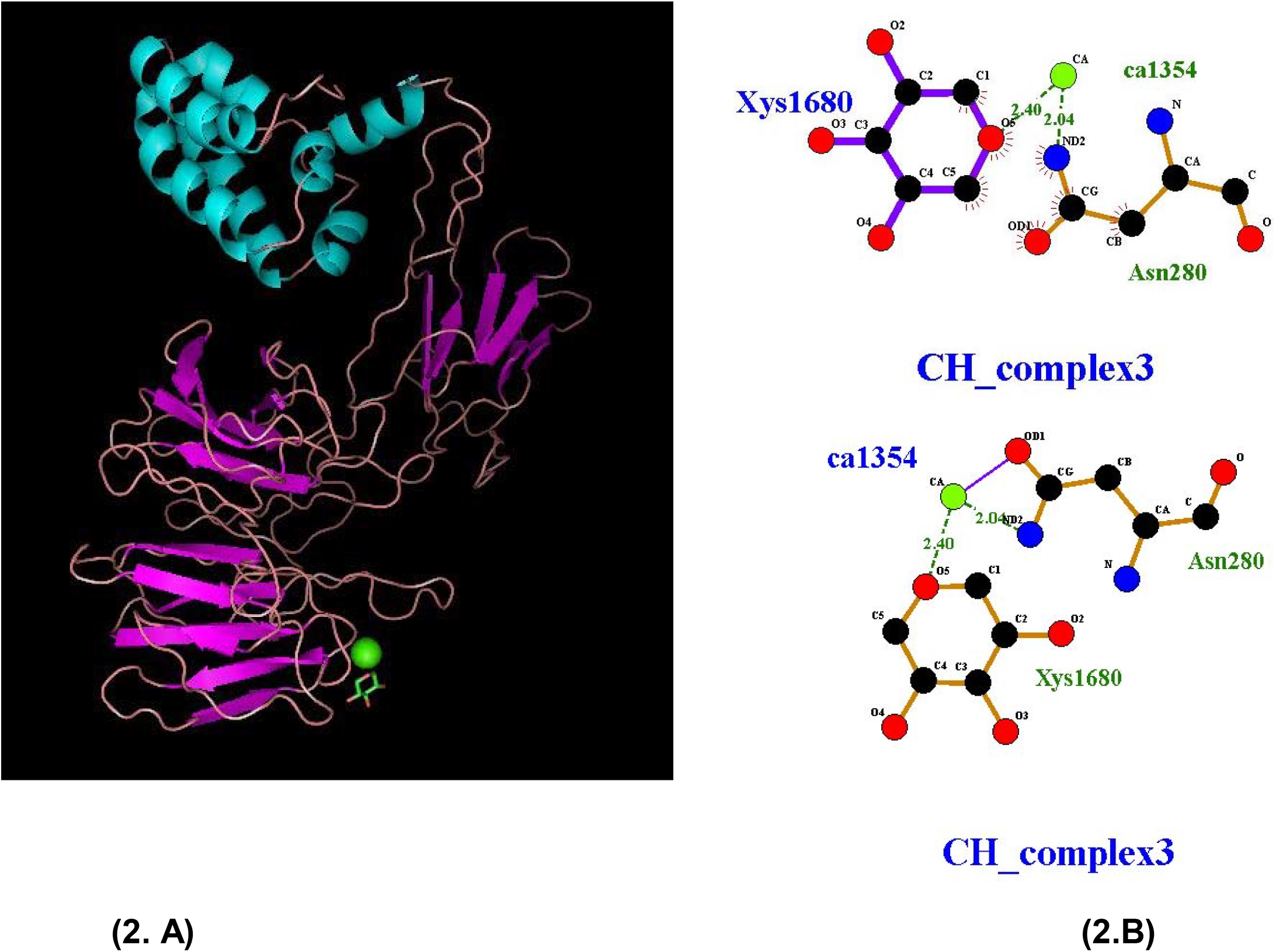
**(A)**Showing the calcium binding site in the beta parallel roll.**(B)** Showing the protein ligand interaction in LigPlus software. [Calcium (ca1354)]

### 3. Docking results shows interaction with drug molecules and molecules involved in apoptosis

String analysis showed there was a e-caprolactam degradation pathway, there are three genes echA8, echA9 and fadB2, showed co-expressed with target protein,(as shown in table 2) investigating its structure and function it was hypothesized that there might be the possibility that the protein was interacting with the different anti-TB drugs. So, based on this finding, the protein-docking study was carried out by using autodockvina and Pymol. A few drugs were selected to dock with the protein model of PE_PGRS20. One of the models of protein interacting with cycloserine is shown in figure 3.A and 3.B. The major interacting residues are THR444 and ASN426. (The list of anti-TB drugs and their binding affinities are shown in table 3.)

**Table 1:** Top 10 EspR binding loci from ChIP-Seq.

| Gene | Position | Gene description | Functional category |
| --- | --- | --- | --- |
| *rv2930* | inside | FadD26, fatty-acid-CoA synthetase, PDIM production | lipid metabolism |
| *rv1490* | overlaps start | Rv1490, probable membrane protein | cell wall and cell processes |
| *rv2929* | overlaps start | Rv2929, PDIM production | conserved hypotheticals |
| *rv1067c* | overlaps start | PE_PGRS family protein, PE_PGRS19 | PE/PPE |
| *rv3616c* | upstream | EspA, ESX-1 secretion-associated protein A | cell wall and cell processes |
| *rv1753c* | inside | PPE family protein, PPE24 | PE/PPE |
| *rv0986* | inside | ATP-binding protein of ABC transporter | cell wall and cell processes |
| *rv1068c* | upstream | PE-PGRS family protein, PE_PGRS20 | PE/PPE |
| *rv3487c* | upstream | LipF, probable esterase/lipase | lipid metabolism |
| *rv0469* | upstream | UmaA, Mycolic methyltransferase A 1 | lipid metabolism |

**Table 2:**
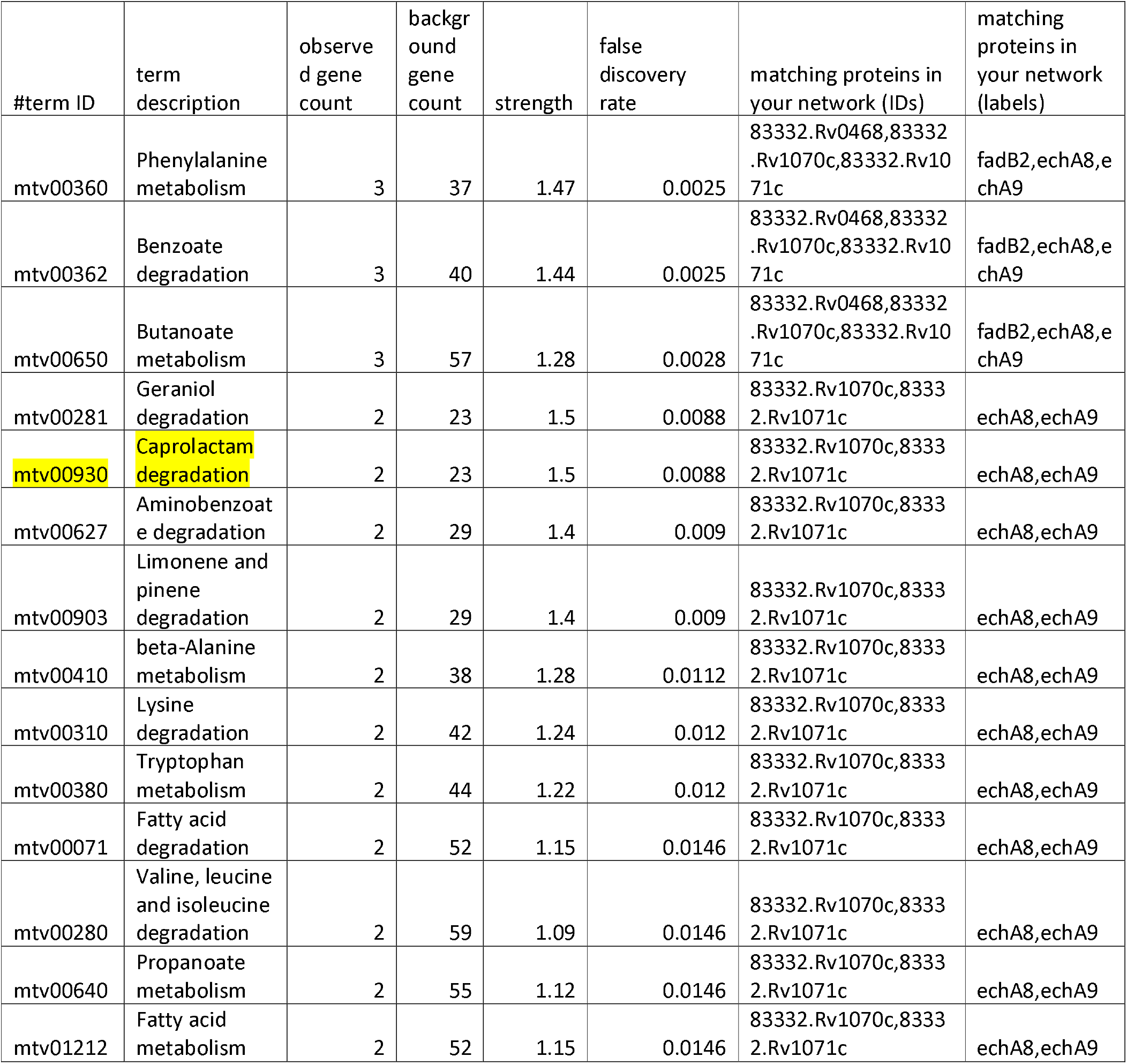
Showing the role of echA8 and echA9 in caprolactam degradation (xenobiotics degradation pathway).

**Table 3:** Docking of target protein with different anti T.B. FDA approved drugs.

| Ligands | PubChem ID | Binding Affinity |
| --- | --- | --- |
| Ethambutol | 14052 | -4.7 kcal/mol |
| Isoniazid | 3767 | -5.4 kcal/mol |
| Pyrazinamide | 1046 | -4.8 kcal/mol |
| D-Cycloserine | 6234 | -4.5 kcal/mol |

**Figure:3.**
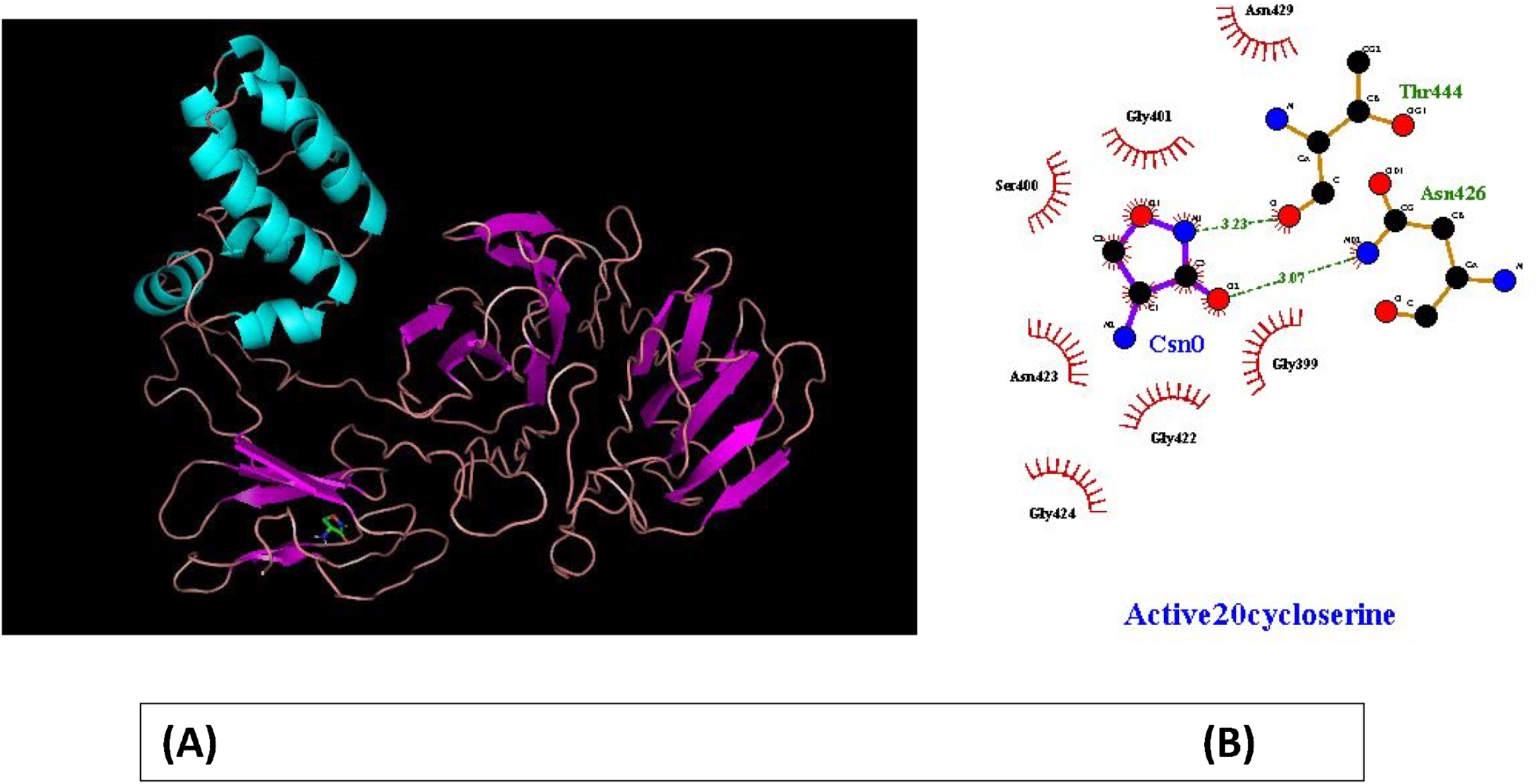
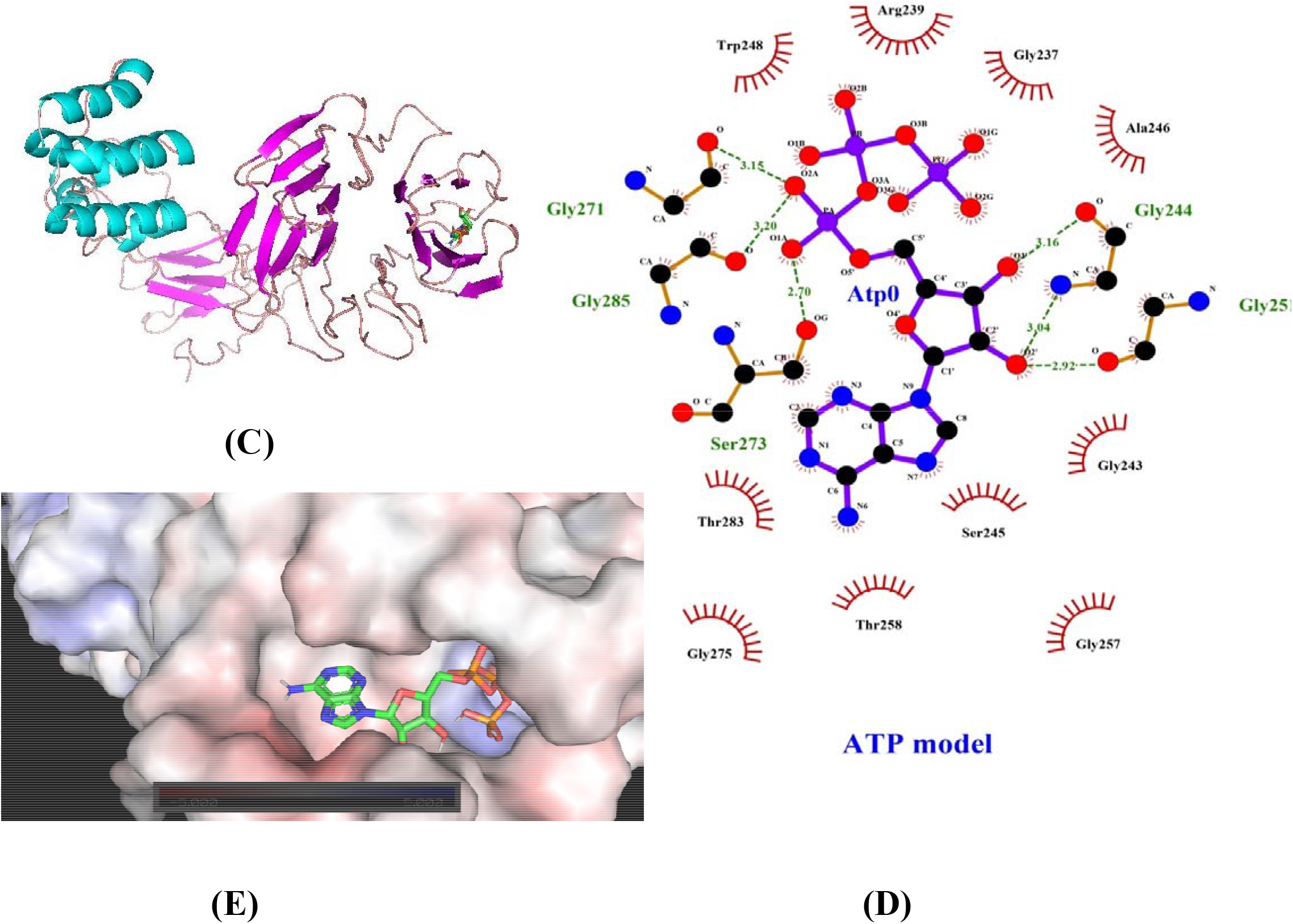
**(A)** cycloserine drug in terminal beta sheets of PE_PGRS20 protein. **3(B)** Diagram showing interacting residues of protein with cycloserine drug.**3(C)** Diagram showing ATP (ligand) in beta-roll of PE_PGRS20.**3(D)** Diagram showing interacting residues of protein with ATP. **3(E)** Binding pocket of ATP in protein.

In addition to that, the protein is involved in the apoptosis of host cells **(GO:0016505)** and has caspase regulator activity.^9^ The quickGO annotation showed that there is binding with ATP(**<u>GO:0005524</u>**).To test this hypothesis we used docking of ATP with the target protein structure. The docking study shows that ATP has binding affinity of −8.9 kcal/mol. The interacting amino acid residues are GLY244, GLY251, GLY271, GLY285, SER273. (Figure 3.C and D). Another docking used FAS molecule as ligand and shows the higher binding affinity (−9.9 kcal/mol) with target protein molecule. NAD molecule can also bind with the target protein molecule with highest binding affinity of −10.7 kcal/mol (see supplementary data).

### 4. Gene Ontology shows possible role of PE_PGRS20 protein in apoptosis of host cell

The DNA sequence of PE_PGRS20 was submitted to the quickGO server. PE_PGRS20 protein function was investigated through homology method with the other matching DNA sequence. This hypothesis was made based on the data, that the target protein is a apoptotic protease activator **(GO:0016505)**, has essential role in apoptosis of host cells **(GO:0006915)** and caspase regulator activity (**GO:0043028)**. Data retrieved from string database which showed that the target protein is involved in the ESX-1 secretion system and takes part in the virulence of *M. tuberculosis* and transcriptionally regulated by EspR protein which is a major virulence factor. More recently, the pro□apoptotic member of the Bcl□2 family, the BH3□only protein Bim was proposed to be involved in the apoptosis induced by virulent *M. tuberculosis*, in a manner that likely involves the ESX□1secretion system^18^.It has been showed that the apoptosis induced by *M. tuberculosis* is abolished in the presence of inhibitors of caspase□9 and caspase□3,suggesting that the capsase□9 dependent formation of an apoptosome structure and the activation of the executioner caspase□3 are required.^19^

### 5. Molecular dynamic simulations and salt bridge analysis

Molecular Dynamic simulations are the tools to identify different energies of the protein system and residue’s trajectory. Hence, Simulation of protein structure can give overall structure stability as shown in parameters (see Table.4) The average value of pressure during the equilibration phase remains around 5.785175 Bar which shows that the protein structure is stable under different pressure. The density value was calculated to be averaging around 1014.284 Kg/m3 which is comparable to the experimental value of water (1000 Kg/m3). The normal values of energies are shown in table 4. This shows that the system is stable over the entire course of simulation (fig. 6.A-D). The changes in residues occur before and after the molecular dynamics simulation were observed as shown in fig.7. ^20^

**Table 4:** Gromacs molecular dynamic parameter for PE_PGRS20 protein shows the stability of protein structure.

| GROMACS molecular dynamics parameters of PE_PGRS20 protein |  |  |
| --- | --- | --- |
| Parameter | Average | Units |
| Potential | -1.8e+06 | KJ/mol |
| Density | 1014.284 | Kg/m<sup>3</sup> |
| Temperature | 299.8955 | Kelvin |
| Pressure | 5.785175 | Bar |

**Figure 4.**
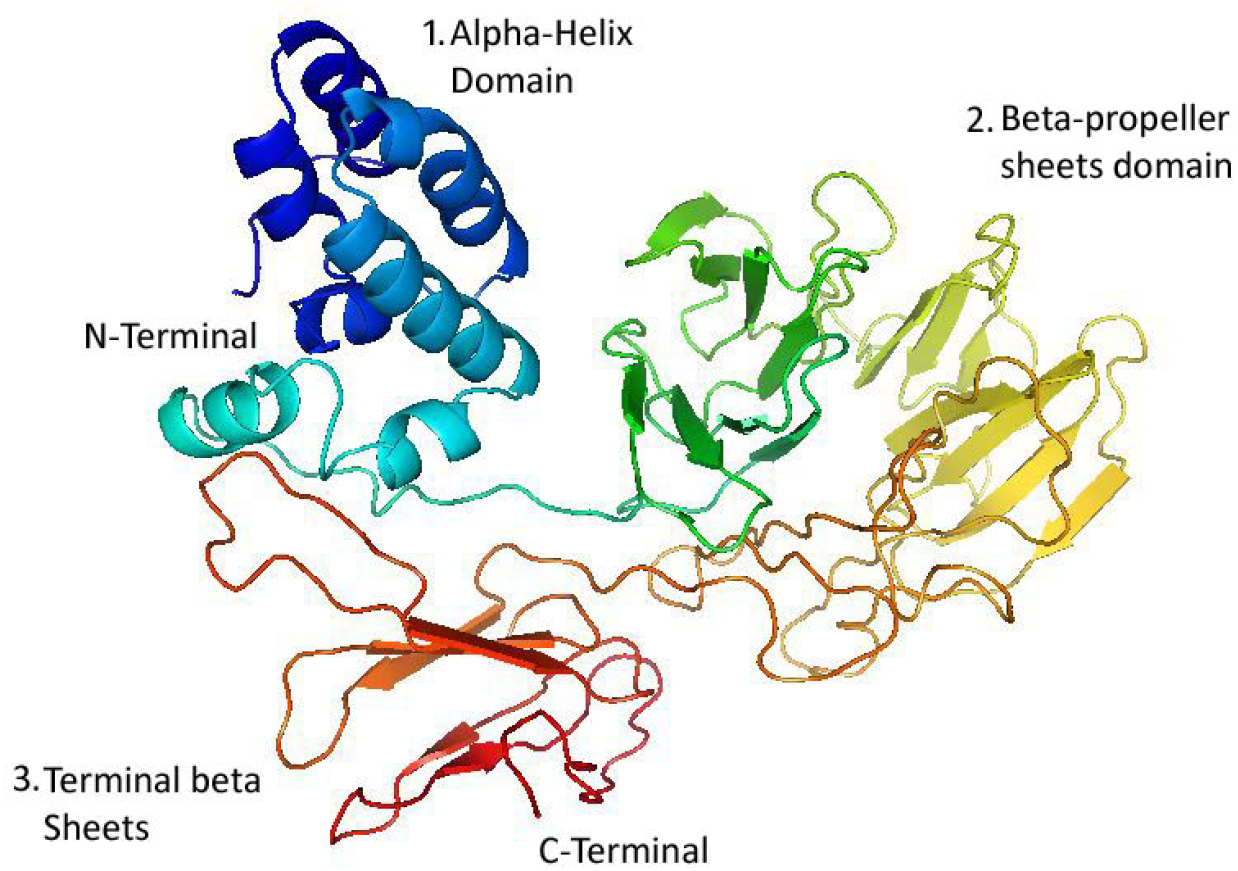
**Structure of PE_PGRS20 protein**

**Figure 4(A).**
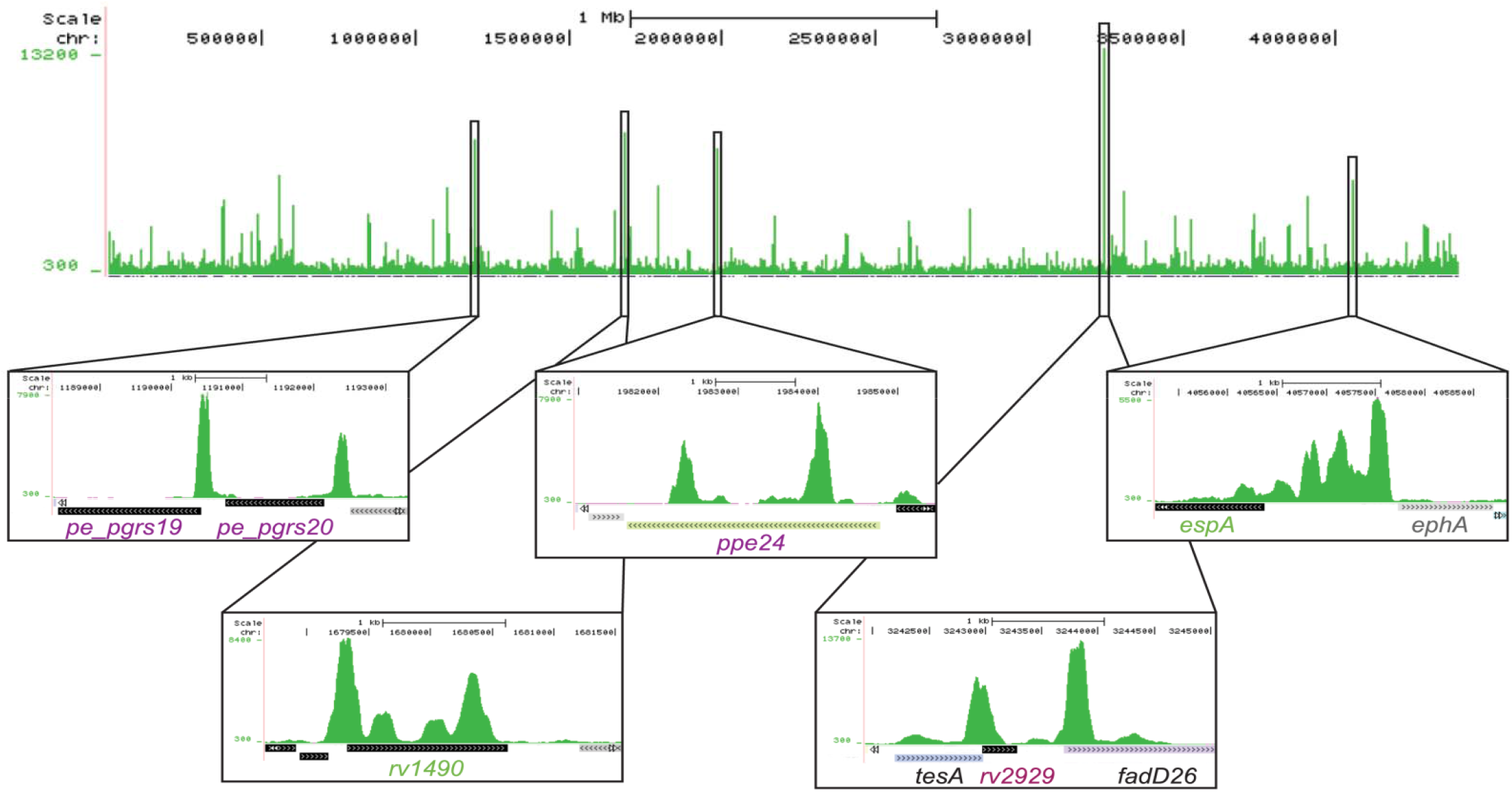
Genome-wide mapping of EspR binding sites. UCSC Genome Browser (http://genome.ucsc.edu) view of EspR binding across the Mtb genome as determined by ChIP-Seq. Peak height (y-axis) indicates the sequencing read depth at each genomic position (x-axis). Inset boxes show major EspR binding sites identified over (from left to right) the pe_pgrs19 and pe_pgrs20 genes, the rv1490 gene, the ppe24 gene, the rv2929 and fadD26 genes from the PDIM/PGL locus and the espACD operon promoter region. doi:10.1371/journal.ppat.1002621.g001^17^

**Fig 4 (B):**
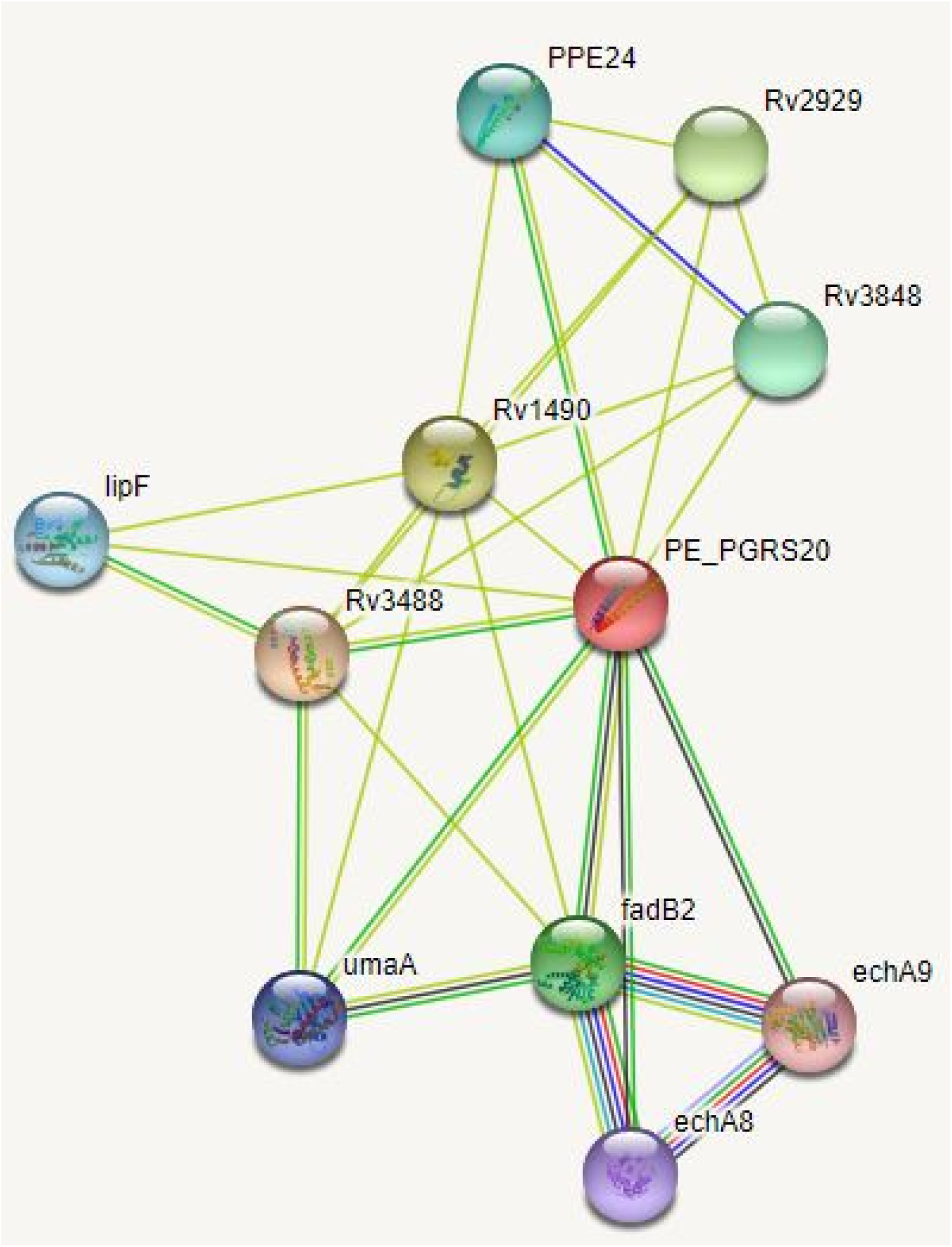
Showing protein co-expression of PE_PGRS20 with fadB2, echA9 and echA8.

**Figure 5.1.**
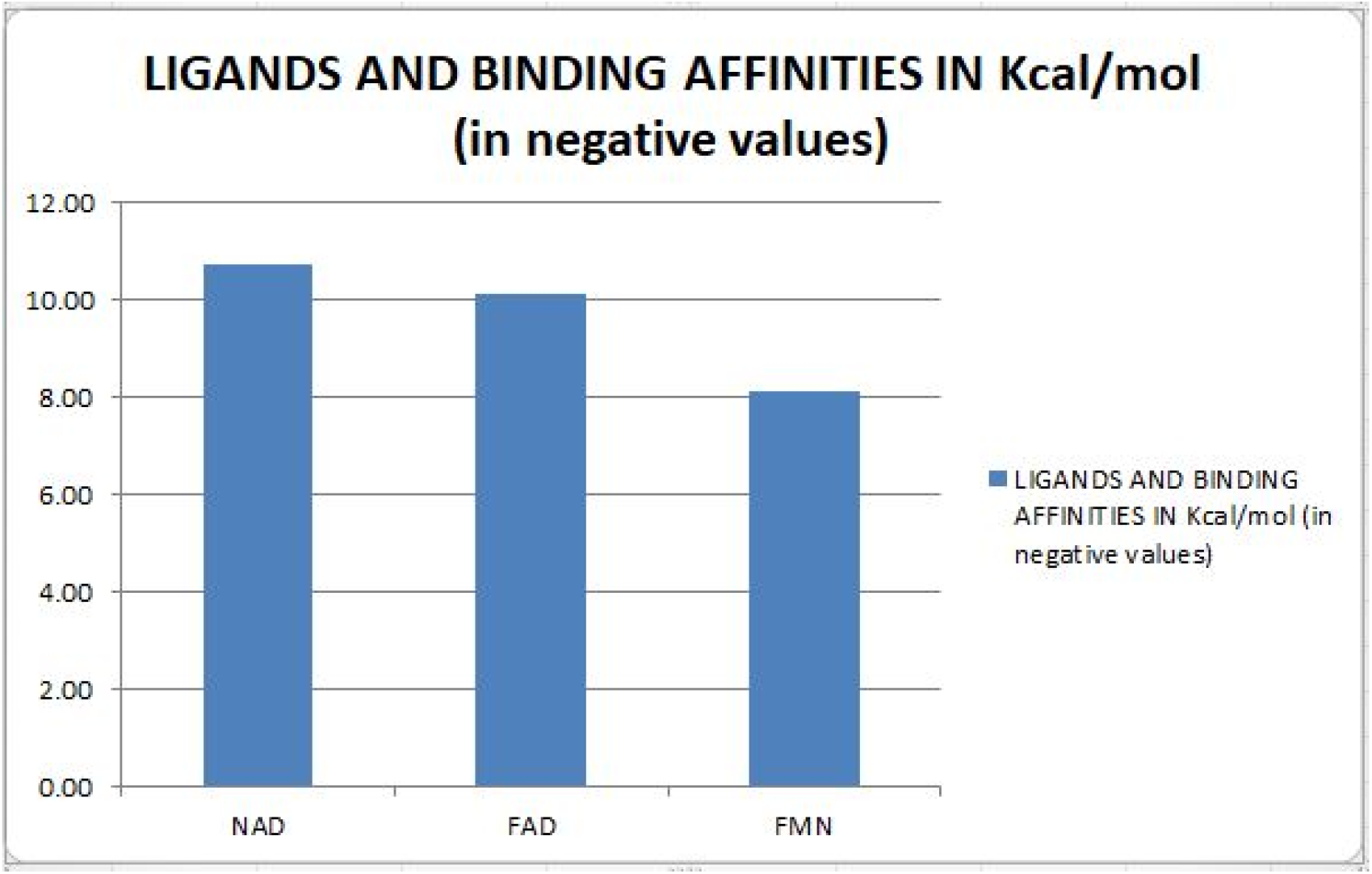
Graph shows binding with NAD, FAD and FMN molecules.

**Figure 5.2.**
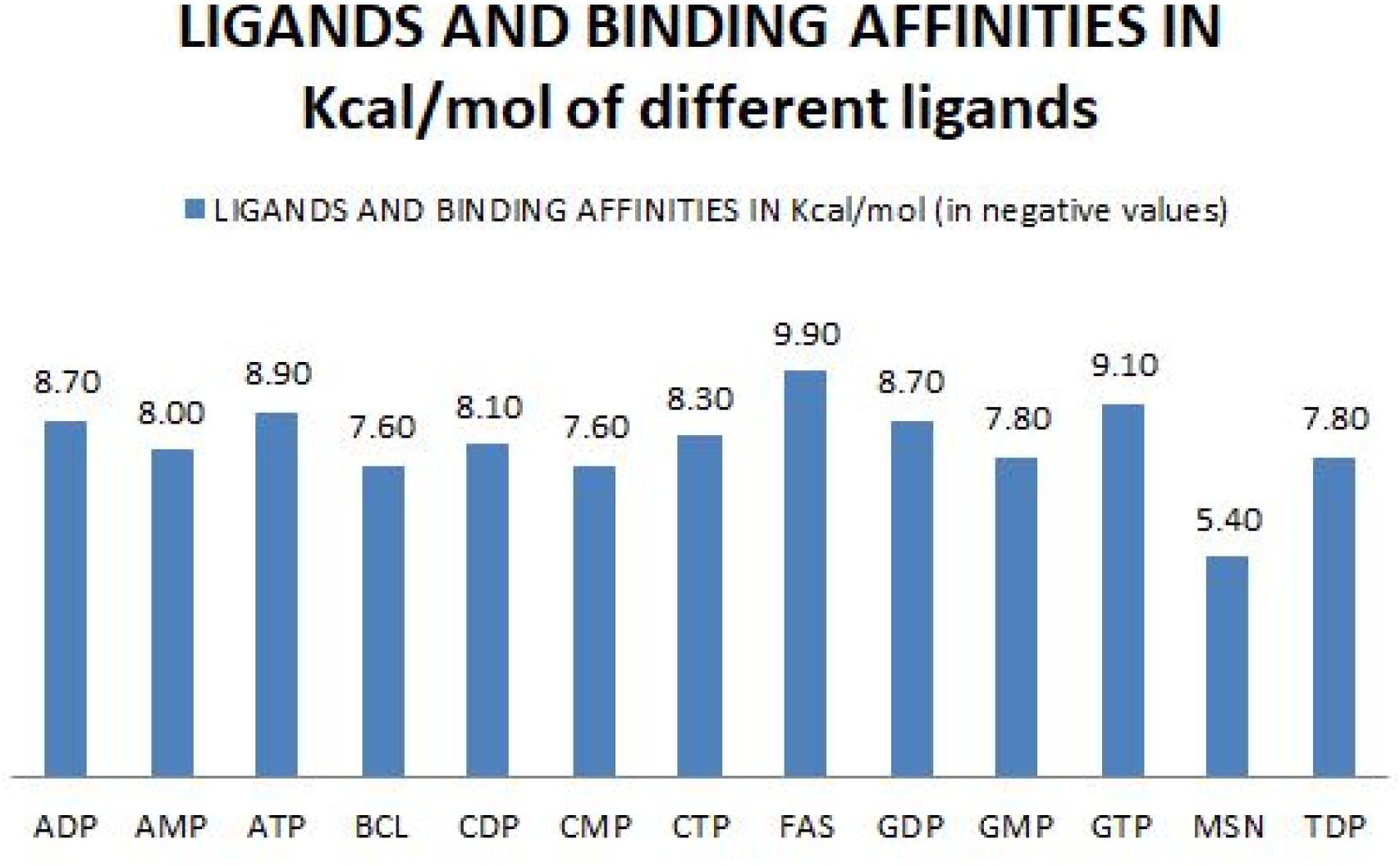
Shows docking with multiple ligands.

**Fig. 6.**
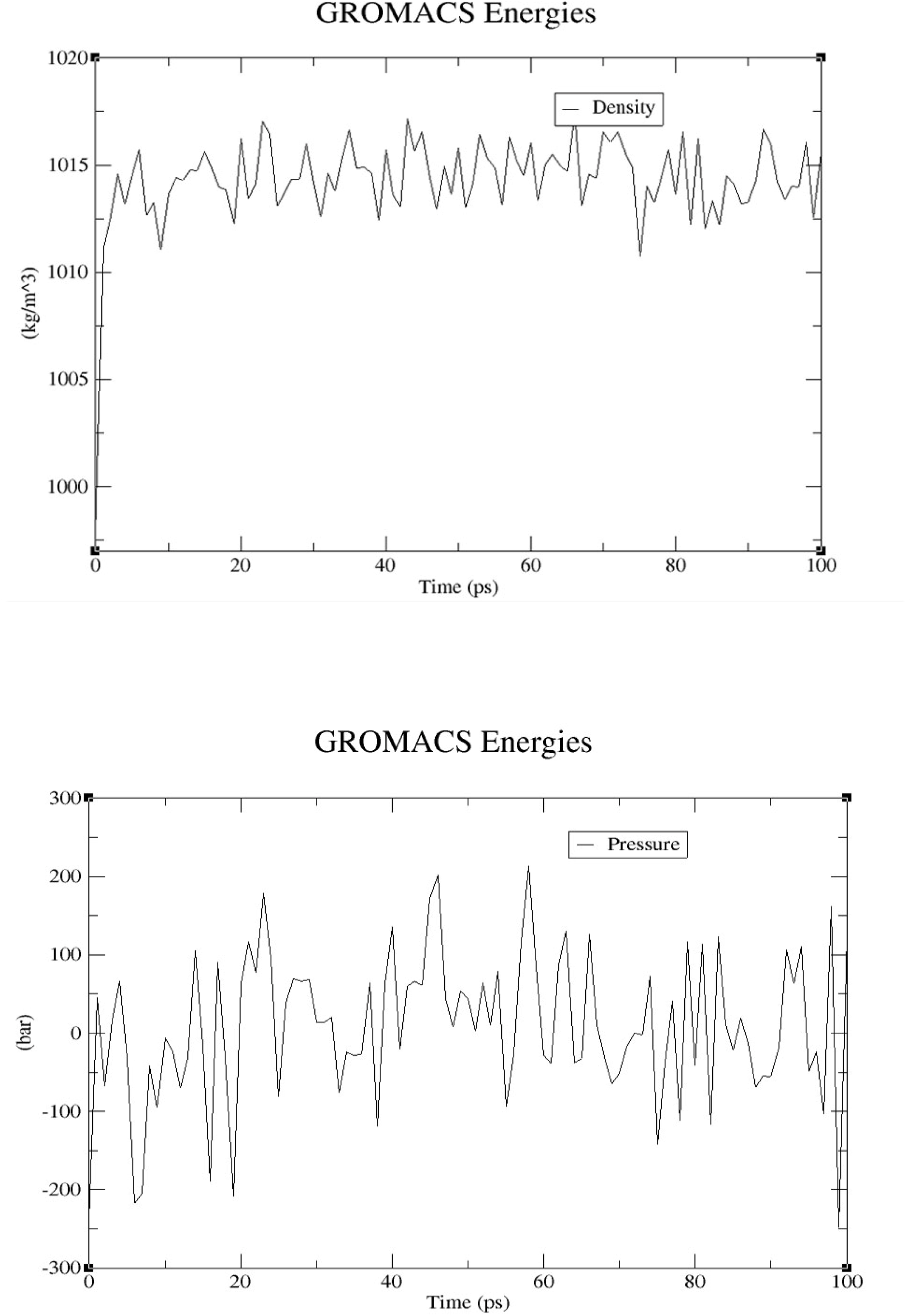
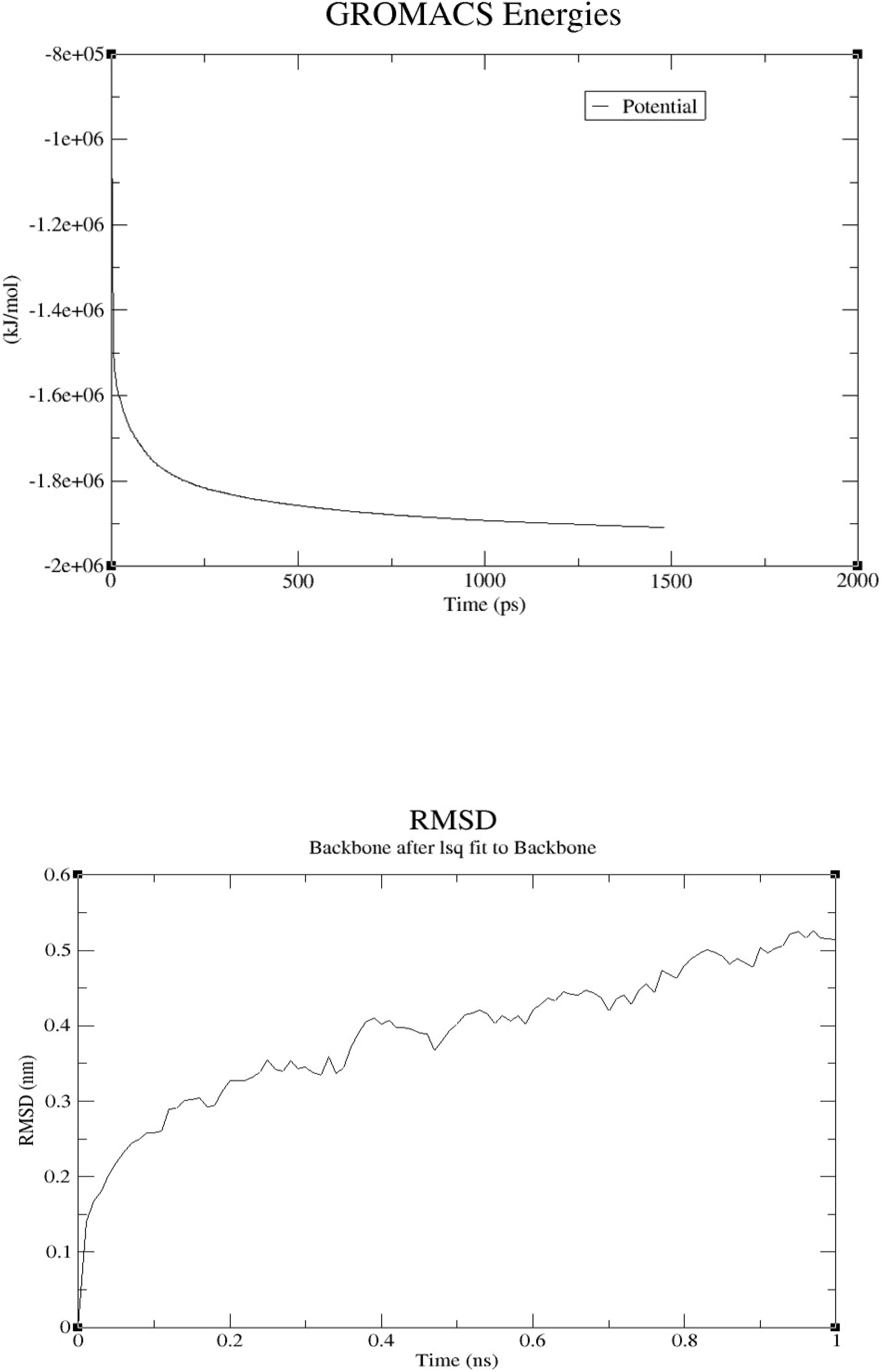
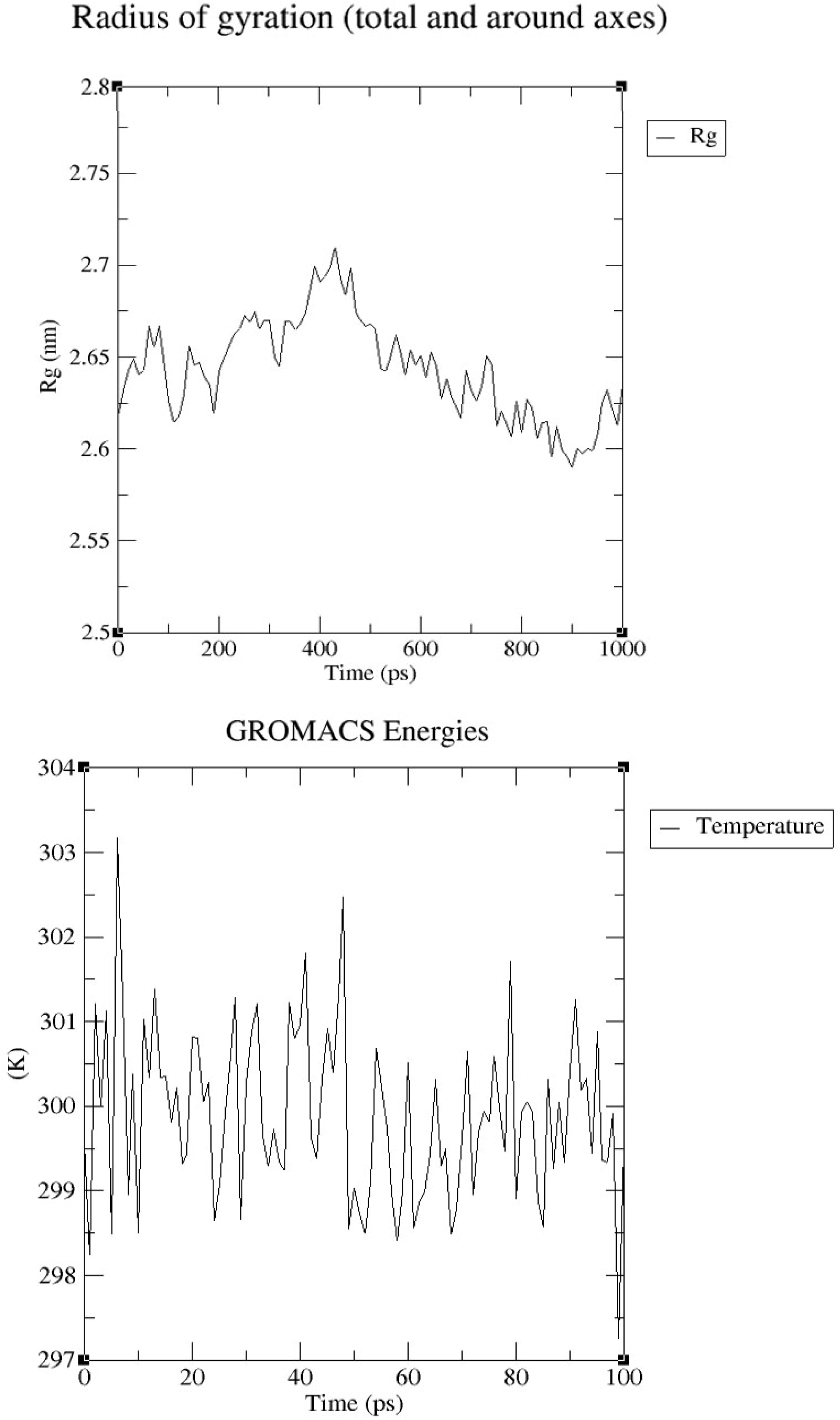
Molecular dynamic simulations over 100 ns showing the structural stability of PE_PGRS20 protein.

There were two salt bridges were identified by VMD tool. HIS58-GLU60 and ASP411-LYS414 (as shown in fig.8.A and 8.B). The salt bridges contribute to the structural stability of protein and maintain the folding-unfolding of protein. They are oppositely charged residues which attracts each other and close to one another.^21^

**Fig. 7:**
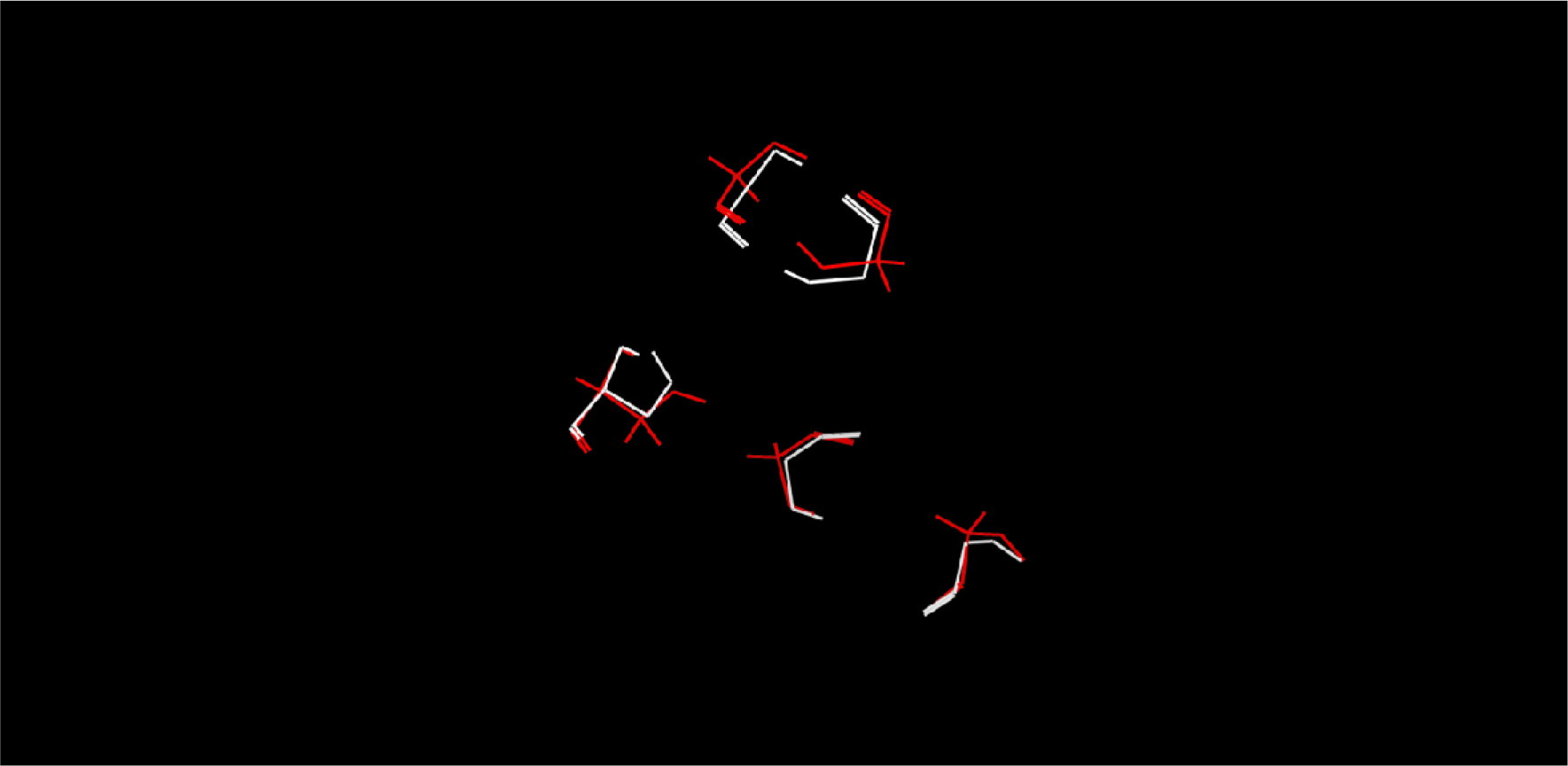
Comparison of structure before molecular dynamic simulation (white) and after molecular dynamic simulation (red).

**Figure 8:**
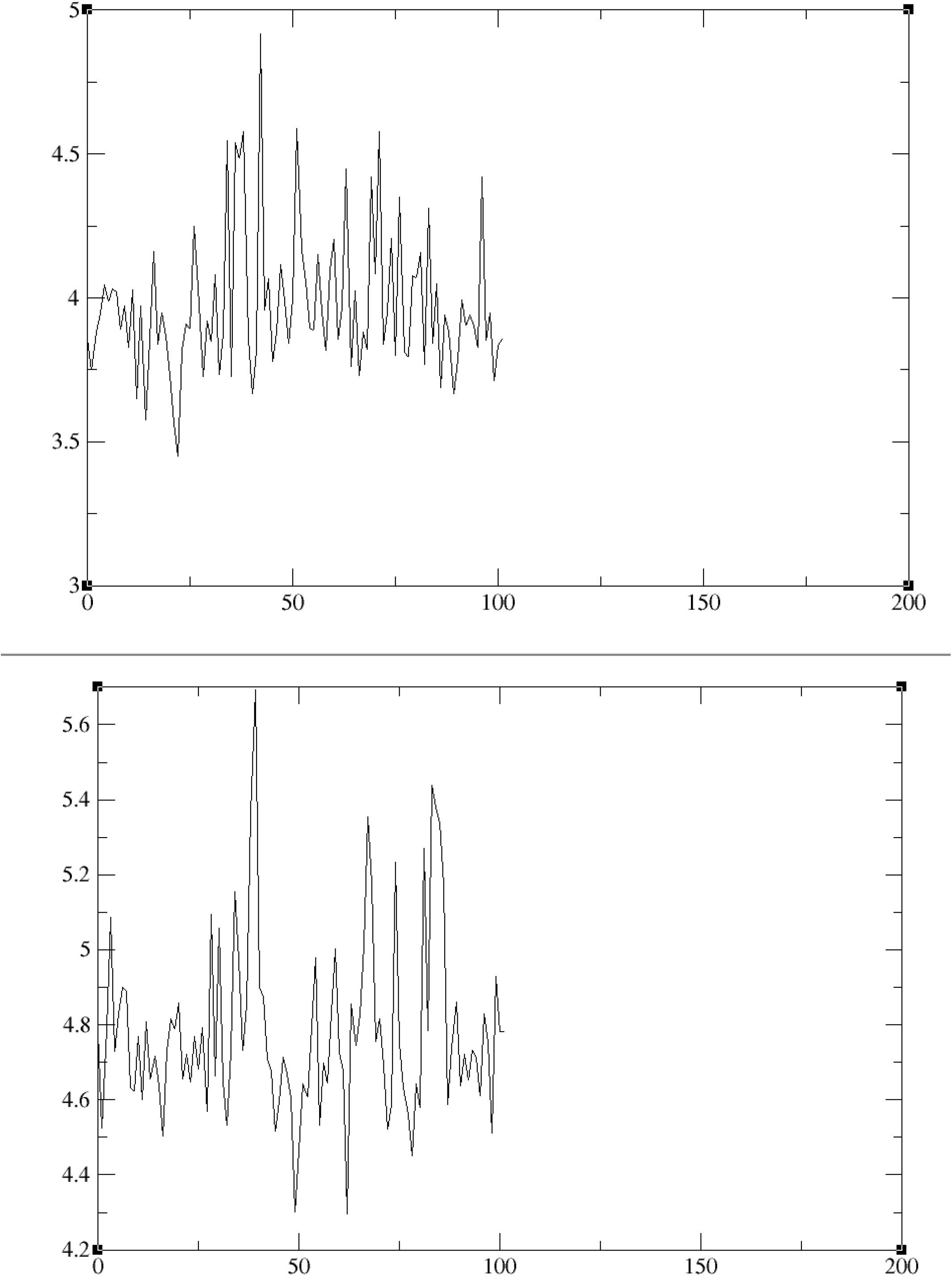
(A) Salt bridge of ASP411-LYS414. (B) Salt bridge of GLU60-HIS58.

The outputs of the MD simulations are trajectories representing snapshots of evolution of the system and appropriate values of time, energy (for example van-der-Waals), applied force, temperature of the system *etc*.^22^

The molecular dynamic simulation is used to observe the structural stability at different temperature, densities and potential etc. It has been observed that the protein structure is stable and to confirm the stability of final modelled structure salt bridge analysis was used.

## Discussion

The final structure was synthesized by using template 4V4L (apoptosome of *Drosophila melanogaster*).The obtained structure was validated by PROCHECK. Ramachandran plot was obtained from the PROCHECK server confirmed the authenticity of the model. The results obtained were further used for protein ligand docking investigations. PE_PGRS proteins are rich in glycine and may have a role in various pathways of *M. tuberculosis*. The PE_PGRS20 protein structure has two catalytic sites (figure. 4). The first catalytic site is present in beta-propeller (Calcium binding site) and the major residues are ASN264, ASN270, SER273, GLY282, GLY296, ASN298, SER301 and ASP304 and second catalytic site is present in terminal beta sheets (Drug binding site) and the major residues is THR444 and ASN426.

The gene PE_PGRS20 of *M. tuberculosis* H37Rv shows binding with EspR protein. EspR protein is responsible for the regulation of ESX-1 system and other Mycobacterial proteins (like PE_PGRS19, PE_PGRS20, FadD26 etc.). EspR is responsible for the virulence of *M. tuberculosis*. EspR functions as a Nucleoid associated protein (NAP) rather than as a specific transcriptional activator of a limited number of genes required for pathogenesis. Diverse functions are encoded by genes where EspR bound upstream and classification by functional category reveals overrepresentation of cell wall/cell processes and the surface-exposed PE/PPE proteins. The detailed mechanism of pathogenesis is still unknown. Among the more than 130 mycobacterial species, only a few have been shown to be able to induce phagosomal rupture and establish contact with the cytosol of the host□phagocytes during infection. All these species harbour a gene cluster encoding an ESX□1 secretion system in their genomes. Phagosomal rupture, which leads to the translocation of bacterial effector proteins into the host cytoplasm and/or to the cytosolic translocation of the pathogen, has important consequences for *M. tuberculosis* recognition by the host and the generation of immune responses. Various proteins of ESX-1 or espACD operon are involved in the lipid synthesis of phthiocerol dimycocerosate (PDIM) and help in the phagosomal rupture and release of mycobacterial protein in the host cell which ultimately leads to the apoptosis of host cell.^23^

## Conclusion

The PE_PGRS proteins are glycine rich protein of unknown structure and function in Mycobacterium tuberculosis H37Rv. Our data suggest that the protein is involved in the apoptosis of the host cell. It has a very important role in ESX-1 secretion system and in the production of lipids (like PDIM) that is necessary for the pathogenesis of *M. tuberculosis* into the host cell. The various proteins are regulated by Nucleoid associated protein (NAP) i.e. EspR.^23^Moreover the molecular dynamic simulations support the structure stability and changes in the active site. The proteins involved in the espACD operon are good targets to understand the mechanism of pathogenesis in mycobacteria. There is no research is reported on the PE_PGRS20 protein yet.

## ACKNOWLEDGEMENT

I would like to pay my humble regard to my project incharge Dr.Anup Kesavan, Assistant Professor, Department of Molecular Biology and Biochemistry, Guru Nanak Dev University, Amritsar for his resolute guidance, unwavering encouragement, motivation, enthusiasm and immense knowledge. I would like to thank him for giving me such opportunity to work on this project.

The above table shows the gene loci at which EspR binds identified by using ChIP-seq

## Notes

### Competing Interest Statement

The authors have declared no competing interest.

## References

1. Brennan MJ, Delogu G. The PE multigene family: a ‘molecular mantra’ for mycobacteria. Trends Microbiol. 2002 May;10(5):246–9. doi: 10.1016/s0966-842x(02)02335-1. PMID: 11973159.

2. Ramakrishnan L, Federspiel NA, Falkow S. Granuloma-specific expression of Mycobacterium virulence proteins from the glycine-rich PE-PGRS family. Science. 2000 May 26;288(5470):1436–9. doi: 10.1126/science.288.5470.1436. PMID: 10827956.

3. Espitia C, Laclette JP, Mondragón-Palomino M, Amador A, Campuzano J, Martens A, Singh M, Cicero R, Zhang Y, Moreno C. The PE-PGRS glycine-rich proteins of Mycobacterium tuberculosis: a new family of fibronectin-binding proteins? Microbiology (Reading). 1999 Dec;145 (Pt 12):3487–3495. doi: 10.1099/00221287-145-12-3487. PMID: 10627046.

4. Dheenadhayalan V, Delogu G, Brennan MJ. Expression of the PE_PGRS 33 protein in Mycobacterium smegmatis triggers necrosis in macrophages and enhanced mycobacterial survival. Microbes Infect. 2006 Jan;8(1):262–72. doi: 10.1016/j.micinf.2005.06.021. Epub 2005 Sep 12. PMID: 16203168.

5. Balaji KN, Goyal G, Narayana Y, Srinivas M, Chaturvedi R, Mohammad S. Apoptosis triggered by Rv1818c, a PE family gene from Mycobacterium tuberculosis is regulated by mitochondrial intermediates in T cells. Microbes Infect. 2007 Mar;9(3):271–81. doi: 10.1016/j.micinf.2006.11.013. Epub 2006 Dec 20. PMID: 17223373.

6. Basu S, Pathak SK, Banerjee A, Pathak S, Bhattacharyya A, Yang Z, Talarico S, Kundu M, Basu J. Execution of macrophage apoptosis by PE_PGRS33 of Mycobacterium tuberculosis is mediated by Toll-like receptor 2-dependent release of tumor necrosis factor-alpha. J Biol Chem. 2007 Jan 12;282(2):1039–50. doi: 10.1074/jbc.M604379200. Epub 2006 Nov 9. PMID: 17095513.

7. Flores J, Espitia C. Differential expression of PE and PE_PGRS genes in Mycobacterium tuberculosis strains. Gene. 2003 Oct 30;318:75–81. doi: 10.1016/s0378-1119(03)00751-0. PMID: 14585500.

8. Prachi P. Singh, Marcela Parra, Nathalie Cadieux and Michael J. Brennan (2008), A comparative study of host response to threeMycobacterium tuberculosis PE_PGRS proteins, Microbiology (2008): 154, 3469–3479.

9. Chen Tian, Xie Jian-ping, Roles of PE_PGRS family in Mycobacterium tuberculosis pathogenesis and novel measures against tuberculosis, Microbial Pathogenesis 49 (2010) 311–314.

10. J Yang, Y Zhang. I-TASSER server: new development for protein structure and function predictions. Nucleic Acids Research, 43: W174–W181 (2015).

11. Dong Xu and Yang Zhang. Improving the Physical Realism and Structural Accuracy of Protein Models by a Two-step Atomic-level Energy Minimization. Biophysical Journal, vol 101, 2525–2534 (2011).

12. Laskowski R A, MacArthur M W, Moss D S, Thornton J M (1993). PROCHECK - a program to check the stereochemical quality of protein structures. J. App. Cryst., 26, 283–291.

13. Trott O, Olson AJ. AutoDock Vina: improving the speed and accuracy of docking with a new scoring function, efficient optimization, and multithreading. J Comput Chem. 2010;31(2):455–461. doi:10.1002/jcc.21334.

14. Laskowski RA, Swindells MB. LigPlot+: multiple ligand-protein interaction diagrams for drug discovery. J Chem Inf Model. 2011 Oct 24;51(10):2778–86. doi: 10.1021/ci200227u. Epub 2011 Oct 5. PMID: 21919503.

15. Nandita Bachhawat and Balvinder Singh, Mycobacterial PE PGRS Proteins Contain Calcium-Binding Motifs with Parallel β-roll Folds (2007), Genomics Proteomics Bioinformatics Vol. 5 No. 3–4(2007).

16. Poulet S and Cole ST 1995 Characterization of the highly abundant polymorphic GC-rich-repetitive sequence (PGRS) present in Mycobacterium tuberculosis. Arch. Microbiol. 163 87–95.

17. LS Meena, Interrelation of Ca^2+^ and PE_PGRS proteins during Mycobacterium tuberculosis pathogenesis, J Biosci(2019) 44:24.

18. Aguiló N, Uranga S, Marinova D, Martín C, Pardo J. Bim is a crucial regulator of apoptosis induced by Mycobacterium tuberculosis. Cell Death Dis. 2014 Jul 17;5(7):e1343. doi: 10.1038/cddis.2014.313. PMID: 25032866; PMCID: PMC4123102.

19. Augenstreich J, Arbues A, Simeone R, et al. ESX□1 and phthiocerol dimycocerosates of Mycobacterium tuberculosis act in concert to cause phagosomal rupture and host cell apoptosis. Cellular Microbiology. 2017;19:e12726.

20. Mukherjee, Tapan. (2016). STRUCTURE PREDICTION AND ASSESSMENT OF BETA-LACTAMASE TEM-1 FROM S. TYPHI USING MOLECULAR DYNAMICS AND SIMULATION STUDIES. International Journal of Recent Scientific Research.

21. Bosshard HR, Marti DN, Jelesarov I. Protein stabilization by salt bridges: concepts, experimental approaches and clarification of some misunderstandings. J Mol Recognit. 2004 Jan-Feb;17(1):1–16. doi: 10.1002/jmr.657. PMID: 14872533.

22. Likhachev IV, Balabaev NK, Galzitskaya OV. Available Instruments for Analyzing Molecular Dynamics Trajectories. Open Biochem J. 2016 Mar 14;10:1–11. doi: 10.2174/1874091X01610010001. PMID: 27053964; PMCID: PMC4797681.

23. Blasco B, Chen JM, Hartkoorn R, Sala C, Uplekar S, et al. (2012) Virulence Regulator EspR of Mycobacterium tuberculosis Is a Nucleoid-Associated Protein. PLoSPathog 8(3): e1002621. doi:10.1371/journal.ppat.1002621

24. Madeira F, Park YM, Lee J, et al. The EMBL-EBI search and sequence analysis toolsAPIs in 2019. Nucleic Acids Research. 2019 Jul;47(W1):W636–W641.

25. Camacho, C., Coulouris, G., Avagyan, V., Ma, N., Papadopoulos, J., Bealer, K., Madden, T.L. BLAST+: architecture and applications. BMC Bioinformatics 10, 421–430 (2009).

26. Remmert, M., Biegert, A., Hauser, A., Söding, J. HHblits: lightning-fast iterative protein sequence searching by HMM-HMM alignment. Nat Methods 9, 173–175 (2012).

27. C Zhang, PL Freddolino, Y Zhang. COFACTOR: improved protein function prediction by combining structure, sequence and protein-protein interaction information. Nucleic Acids Research, 45: W291–W299, 2017.

28. Sigrist CJ, Cerutti L, de Castro E, Langendijk-Genevaux PS, Bulliard V, Bairoch A, Hulo N. PROSITE, a protein domain database for functional characterization and annotation. 1. Nucleic Acids Res. 2010; 38(Database issue):D161–6.

